# Design of a self-regulating mRNA gene circuit

**DOI:** 10.1101/2024.04.23.590729

**Authors:** Eric C. Dykeman

## Abstract

Protein expression from mRNA *in vivo* is predominately controlled via regulatory feedback mechanisms that adjust the level of mRNA transcription. However, for positive sense single-stranded RNA viruses, protein expression is often controlled via secondary structural elements, such as internal ribosomal entry sites, that are encoded within the mRNA. The self-regulation of mRNA translation observed in this class of viruses suggests that it may be possible to design mRNAs that self-regulate their protein expression, enabling the creation of mRNAs for vaccines and other synthetic biology applications where protein levels in the cell can be tightly controlled without feedback to a transcriptional mechanism. As a proof of concept, I design a polycistronic mRNA based on bacteriophage MS2, where the upstream gene is capable of repressing synthesis of the downstream gene. Using a computational tool that simulates ribosome kinetics and the co-translational folding of the mRNA in response, I show that mutations to the mRNA can be identified which enhance the efficiency of the translation and the repression of the downstream gene. The results of this study open up the possibility of designing bespoke mRNA gene circuits in which the amount of protein synthesised in cells are self-regulated for therapeutic or antigenic purposes.

## Introduction

Messenger RNA (mRNA) based therapeutics and vaccines present a promising new platform for the delivery of immunogenic compounds for vaccination or therapeutic proteins for disease treatment and management^1–4^. While there was recently great success with the rapid development of an mRNA based vaccine for SARS-Cov-2, obstacles still remain to the development of mRNA based therapies, which typically require much higher levels of the therapeutic proteins to be delivered to the target cell^1^. This has lead to the development of strategies such as self-amplifying mRNAs^5–7^, where replicase genes are also supplied on the mRNA, allowing it to undergo replication in the cell. However, it is conceivable that there may exist situations in mRNA based pharmacology where protein production in the cell needs to be tightly controlled as opposed to a continuous production of the protein in the cell to extreme levels. Such considerations highlight the need to develop a wide range of strategies for controlling protein expression in mRNA vaccines and therapeutics.

Techniques to regulate gene expression on mRNAs at the translational level have traditionally focused on the use of non-coding RNAs or riboswitches to engineer regulatory mechanisms that can either repress or promote gene expression from the mRNA. For example, trans-acting small RNA translational activators, termed toehold switches^8^, are a class of RNA based regulators of translation that can be designed to interact with mRNAs to promote exposure of the ribosome binding site (RBS), directly controlling protein expression. For example, Wang and Simmel have recently demonstrated that small RNAs can be used to both activate and repress translation by sequestering either anti-anti-RBS or anti-RBS sequences^9^. While Wang and Simmel have used toehold switches to regulate genes on an mRNA, some groups have used toehold based methods to design more complicated mRNA based logic circuits^10^, where expression is triggered by the presence of miRNAs. Attachment of these miRNAs to the mRNA expose the RBS, providing potential cell specific expression of the mRNA. In addition to toehold based methods, Hong *et al*.^11^ also employed the use of small transcriptional activating RNA (STAR)^12–14^, which target transcriptional activation by the RNA polymerase, to construct an incoherent feed-forward loop to control gene expression. Both the use of STAR and toehold switches rely on external small RNAs to exert regulatory control on protein expression, which may be difficult to employ broadly in mRNA based therapeutics.

In contrast to regulatory control of translation using small RNAs, bacteria and viruses provide many alternative examples of how RNA secondary structure within the mRNA, and its interactions with metabolites and proteins, can provide direct regulatory control of gene expression at the translational level. For example, the small positive sense single-stranded RNA (ssRNA) virus bacteriophage MS2 regulates the synthesis of its RNA dependant RNA polymerase (RdRp) gene via a feedback mechanism that tightly controls the amount of RdRp produced in relation to the amount of coat protein present. Specifically, coat protein dimers bind to a 19 nucleotide hairpin encompassing the start codon for the RdRp gene with nanomolar affinity. Thus, as coat protein concentration increases, binding of coat protein to the hairpin blocks further ribosome initiations on the RdRp gene^15^. Similar to phage MS2, bacterial mRNAs also have similar feedback mechanisms which regulate protein expression from the mRNA transcript and monitor free levels of the protein in the cell. For example, in the L11-L1 polycistronic mRNA in *E. coli*, as excess amounts of the large ribosomal protein L1 protein accumulates in cell, the L1 protein binds to the RBS of the L11 protein in the mRNA and slows expression of both proteins^16^. These examples provide an alternative strategy to that of trans-acting small RNAs for the purpose of regulating protein production in mRNAs, where feedback to a transcriptional control mechanism or control via small trans-acting RNAs is not feasible.

In this work, I report on a proof of concept application where I have designed a polycystronic mRNA in which the protein production is self-regulated. Specifically, I have re-purposed the MS2 translation repression and coupling mechanism, which functions to regulate the level of the RdRp protein produced, to instead regulate the levels of the 19 kDa NanoLuc luciferase (NLuc) protein. The resulting polycistronic mRNA contains separate genes for the coat and NLuc proteins, with the NLuc protein being translationally coupled and repressed by the MS2 coat protein. Through the use of my computational tool which simulates the kinetics of ribosome translation and mRNA co-translation folding due to ribosome movements on the mRNA^17^, I am able to propose a series of mutations which enhance overall protein expression and the ability of MS2 coat protein to repress expression of NLuc via the binding of coat protein to a TR hairpin. Interestingly, the results also reveal the importance of alternative RNA secondary structures in the RBS and how these can kinetically compete to alter gene expression, providing important insights into the impact of RNA secondary structure on translation efficiency.

## Methods

### 0.1 Reagents

Synthetic oligodeoxynucleotides were custom ordered from Eurofins Genomics and the pET-21a(+) cloning vector was supplied by Novagen (#69770). Restriction enzymes were obtained from Thermofisher (#FD0083, #FD0094). NanoGlo assay reagent was obtained from Promega (#N1110). Restriction digestion clean-up and plasmid isolation was performed using Thermofisher GeneJet PCR clean-up and miniprep kits (#K0701, #K0502). Mutagenesis of the plasmids was done using the NEB-Q5 mutagenesis kit New England Biolabs (#E0554S). All commercial enzymes and kits were used following their provided protocol and the manufacturers provided buffer(s) unless otherwise stated.

### 0.2 Biological Resources

Competent *E. coli* DH5*α* cells for cloning were obtained from Thermofisher (#EC0112) and competent *E. coli* BL21 (DE3) cells for expression were obtained from New England Biolabs (#C2427H).

### 0.3 Plasmid preparation and mutagenesis

Two DNA fragments encoding the CTnano-cnt and CTnano mRNAs were synthesised (GeneStrands, Eurofins Genomics, Cologne, Germany) with each containing the unique restriction sites BglII and Bpu1102. The CTnano-cnt mRNA sequence is based on bacteriophage MS2 coat gene (Genebank accession code NC001417) and a NanoLuc luciferase gene which has been codon optimised for expression in *E. coli* (Genebank accession code MN834152). The CTnano mRNA sequence is a synonymously re-coded version of CTnano-cnt designed for enhanced expression of NLuc. The exact sequences synthesised are given in supplementary data. These sequences were separately cloned into the pET-21a(+) vector (Novagen) using the restriction sites BglII/Bpu1102 resulting in the plasmids pET-CTnano and pET-CTnano-cnt. Production of the mRNAs in these plasmids are under the control of a T7 promoter and are terminated by a T7 terminator hairpin (c.f. Supplementary Figure 1). Following creation of these plasmids, three mutations (N55D, T19*, and S37*) where separately introduced to the wild-type MS2 coat protein present in these mRNA constructs using the NEB-Q5 mutagenesis kit (New England Biolabs). The primer sets and PCR settings used are listed in supplementary information. Mutations were subsequently verified by Sanger sequencing of the plasmids (Eurofins Genomics) and all plasmids were grown and isolated using GeneJet plasmid miniprep (Thermofisher) from *E. coli* DH5*α* cells (Thermofisher).

### 0.4 NanoLuc luciferase luminescence assay

New England Biolabs *E. coli* BL21 (DE3) cells containing one of the 8 plasmids used in this study were grown at 37° C in 5 ml of LB broth supplemented with 100 *μ*g/ml ampicillin. When cells reached an optical density *A*_600_ ≈ 0.6, expression of the mRNA was induced using 0.1 mM isopropyl *β* -D-thiogalacoside (IPTG). After 30 min of incubation at 37° C, the final *A*_600_ of the culture was measured and 20 *μ*l of culture was diluted with 80*μ*l of fresh LB and added to 100 *μ*l of NanoGlo luciferase assay (Promega) consisting of NanoGlo lysis buffer supplemented with the provided furimazine substrate at a v/v ratio of 50:1. Measurements of the luminescence were performed on a BMG Labtech Flowstar OPTIMA 96 well plate reader with emission filter set to 460nm at time points of approximately 5, 15, 30, 45, and 60 min after addition of the NanoGlo assay reagent. Detection limits of the instrument were determined to range between 2.1 *×* 10^2^ and 2 *×* 10^6^ (Arb. U.). Measurements were performed in triplicate using three different samples of culture and luminescence time courses were fit to the equation *L*(*t*) = *a* + *b/*(1 + *exp*(−*c*(*t* −*d*))), where *a, b, c* and *d* are constants as detailed in Supplementary Figures 5 and 6.

### 0.5 Computational prediction of mRNA structure

To compute putative secondary structures of both the CTnano-cnt and CTnano mRNAs, I assume that the sequence upstream of the TR stem-loop folds both co-transcriptionally and co-translationally into the native structure by the time the polymerase or ribosome reaches the TR stem-loop. Following this assumption, I enforce that the long-distance Minju interaction is in place and fix the secondary structure of the mRNA upstream of the TR stem-loop to that of the native structure of the MS2 coat gene, which has been determined experimentally using enzymatic probing and phylogenetic analysis^18,19^. The secondary structure of the remaining downstream sequence is determined from the minimum thermodynamic free energy fold computed using Turner energy rules^20^. The resulting structures are shown in Supplementary Figures 2 and 3.

### 0.6 Computational design of CTnano mRNA for enhanced NLuc expression

Potential synonymous mutations to the NanoLuc luciferase gene which stabilise the TR hairpin and other long distance interactions are identified and tested using a three pronged approach. First, the minimum free energy structure of the mutated sequence is computed using standard mRNA structure prediction algorithms. Second, the mutated sequence is folded for *t* = 1 second of time kinetically using KFOLD^21^ starting from a single-stranded state. Finally, the mutated sequence is tested for its ability to return the NLuc ribosome binding site back to its original structure during co-translational folding in the presence of ribosomes using my ribosome/folding kinetics model which simulates mRNA co-translational folding kinetics due to ribosome movement over the mRNA^17^. Satisfying these three tests insures that; (a) kinetic folding of the sequence is “funnelled” into the thermodynamic minimum free energy structure and, (b) that co-translational folding of the sequence does not trap any structures (particularly the ribosome binding site) into mis-folded states.

The overall procedure used to identify the CTnano mRNA sequence is as follows. First, a set of mutations are identified which forces the desired structure to have the thermodynamic minimum free energy. Next the sequence is tested using the ribosome folding model^17^ for co-translational folding of the mRNA and its ability to return the NLuc ribosome binding site back to its original structure. Subsequent mutations are then made which improve the co-translational folding properties of the NLuc ribosome binding site while also testing that these mutations do not alter the minimum free energy structure. Finally, KFOLD^21^ is used to check that the sequence folds kinetically into the thermodynamic minimum free energy structure starting from a single-stranded state. Although a total of 77 mutations to the CTnano-cnt were required, it should be noted that many of these were chosen to stabilise structures downstream from the NLuc ribosome binding site. This was done as a precaution to avoid the possibility that long distance interactions with these areas form with either the coat region (first 478 nucleotides of the CTnano mRNA sequence) or the NLuc ribosome binding site.

### 0.7 Computational simulation of gene expression

A variety of computational tools exist for the estimation of gene expression which take into account the secondary structure of the mRNA around the RBS and any potential interactions of the 30S subunit with the Shine-Dalgarno sequence. These include the RBS Designer^22^, the RBS Calculator^23,24^, and the UTR Designer^25^. While each of these can provide estimation of gene expression in a prokaryotic system, they are not capable of predicting gene experssion on mRNAs containing the two key features of interest in this study, specifically translational coupling and repression. For this reason, computational prediction of coat protein and nano luciferase expression of the CTNano-cnt and CTnano mRNAs where simulated using my ribosome/folding kinetics model^17^. This model is capable of predicting gene experssion in mRNAs with translational coupling/repression since it simulates the movement of ribosomes and the resulting co-translational folding of the mRNA of interest in addition to simulating the entire translational process that would be occurring in the cellular ‘background’ of an exponentially growing *E. Coli* cell. For example, in an *E. Coli* cell with doubling time 60 min, my model accounts for translation from 15000 ribosomes on ≈ 1200 cellular mRNAs and includes all known kinetic steps in the translation process for initiation, elongation, and termination. In addition to taking into account the competition between mRNAs for available free ribosomes, the model also considers the competition of tRNAs for the A-site during decoding, and the recharging of ternary complex by Ef-Ts. Finally, the model also includes the ability of coat proteins to bind to TR-like stem loops in the mRNA and models the competition between ribosomes and coat proteins for the TR stem-loop. Modelling this competition is critical to correctly calculating when the Nano-luc gene will be fully repressed and will be dependent on how the concentration of coat protein increases over time. Further details on the parameter settings, the number of cellular mRNAs, and the cell growth rate, can be found in^17^.

## Results

### Design of a synthetic polycistronic mRNA gene circuit based on bacteriophage MS2

The bacteriophage MS2 regulates the synthesis of its RNA dependent RNA polymerase (RdRp) gene via a combination of two mechanisms, translational coupling and translational repression. In the coupling mechanism, translation of the RdRp gene is dependent on translation of the upstream coat protein gene. Work by Van Duin and colleagues demonstrated that disruption of a long distance RNA secondary structure interaction, the MinJu sequence^26^, by ribosomal movement over the viral mRNA results in the opening of the ribosomal binding site (RBS) for the RdRp gene (c.f. Figure 1a and 1b). In the translational repression mechanism, translation of the RdRp gene is blocked by the binding of a coat protein dimer to the 19 nucleotide translational repressor (TR) stem-loop, which encompasses the start codon for the RdRp gene (Figure 1b). Thus, the overall mRNA secondary structure of the MS2 coat and RdRp genes, as illustrated in Figure 1, enables the synthesis of the RdRp protein to be regulated via feedback from the overall coat protein dimer concentration in the cell.

**Figure 1.**
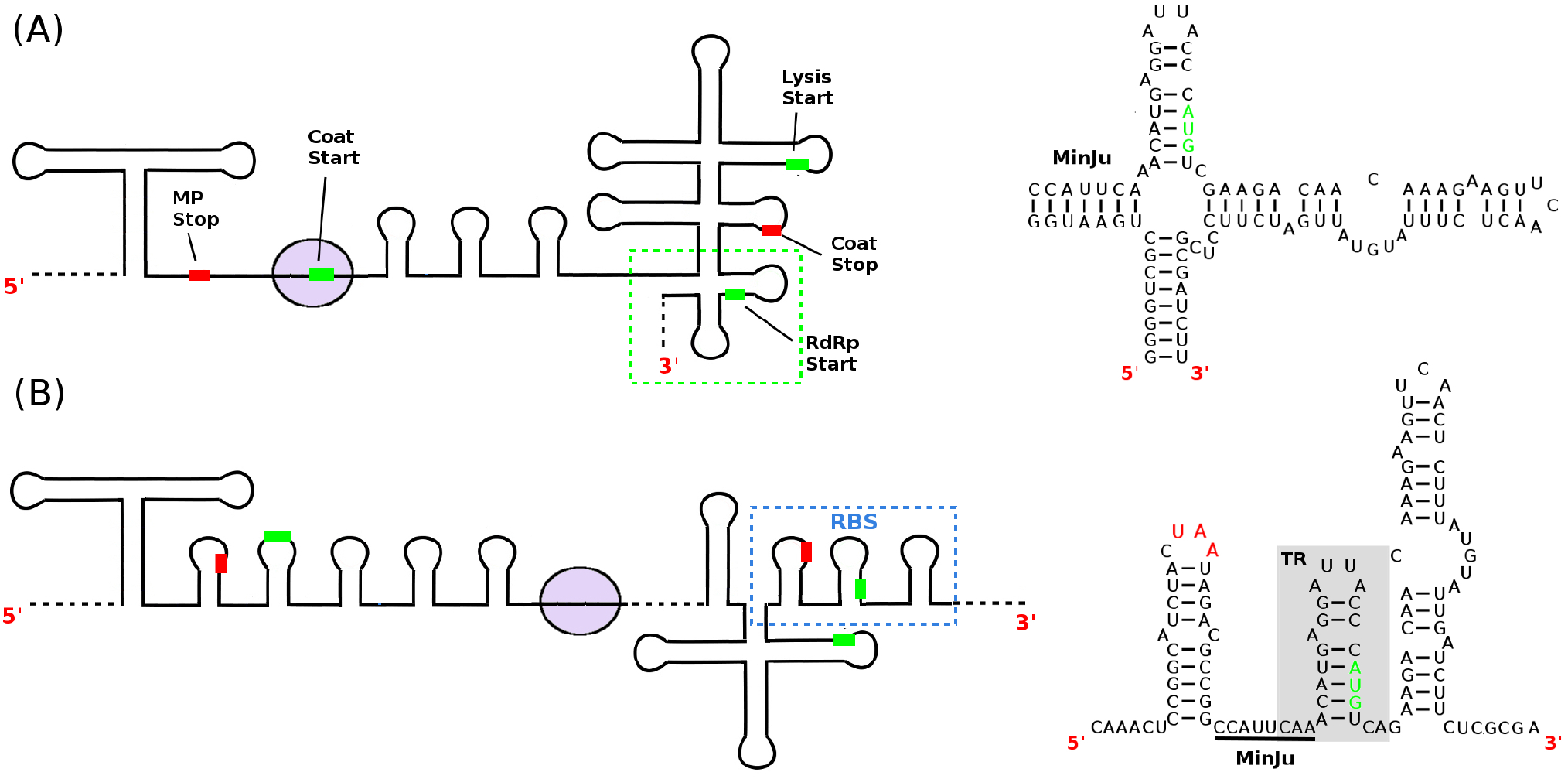
Translational coupling and repression of the RNA dependent RNA polymerase gene in Bacteriophage MS2. (A) Cartoon diagram of a section of the secondary structure of the MS2 viral mRNA (nucleotides 1132-1813)^18,19^ during ribosome initiation on the coat start codon. Approximate locations of the start codons for the coat, lysis, and RdRp genes are labelled by green bars, while stop codons for maturation protein (MP) and coat protein are shown with red bars. The nucleotide sequence of the RdRp ribosome binding site (green dashed box) is shown on the right hand side along with the MinJu long-distance interaction. (B) During ribosome synthesis of the coat protein, the MinJu interaction is disrupted exposing the RdRp ribosome binding site to ribosomes (blue dashed box) with nucleotide sequence shown to the right. When high concentrations of coat protein dimers are present in the cell, they bind to the TR hairpin blocking subsequent ribosome initiations.

To construct a polycistronic mRNA where the NLuc protein is translationally repressed/coupled to the MS2 coat protein, one could simply take the 3’ end of the MS2 genome encompassing coat, lysis and RdRp genes and replace the RdRp gene with a sequence encoding the NLuc protein (as depicted in Figure 2). However, I hypothesise that the overall secondary structure of the NLuc RBS, and its response to co-translational folding, will have substantial effects on the expression of the protein, thus making the choice of codons of the NanoLuc luciferase gene critical to efficient protein expression.

**Figure 2.**
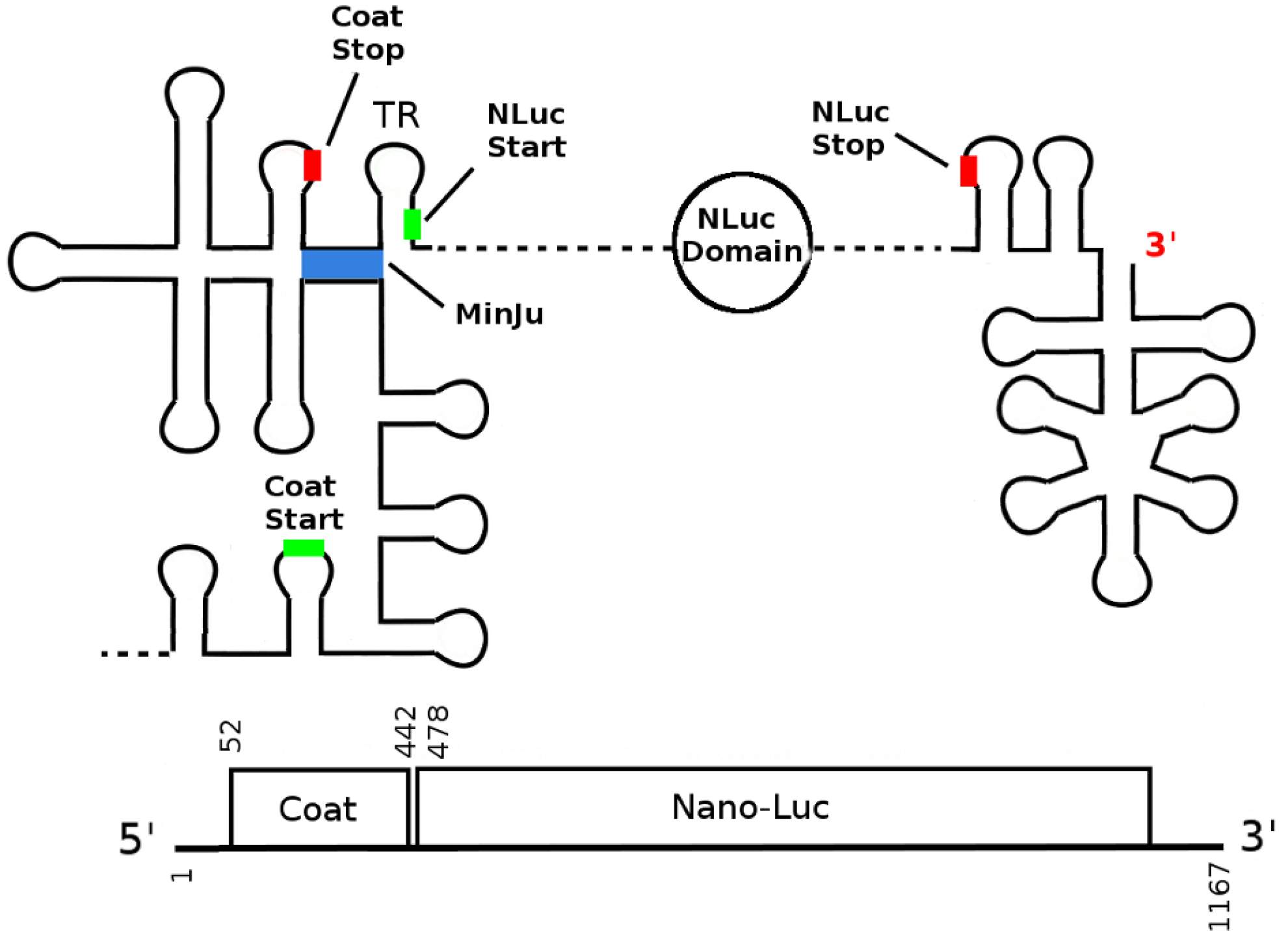
Design of the CTnano-cnt and CTnano mRNAs based on the Bacteriophage MS2 Coat/RdRp gene fragment. The 3’ end of the MS2 viral mRNA (nucleotides 1284-3569) containing the coat, lysis, and RdRp genes is used as a template for construction of the CTnano-cnt and CTnano mRNAs. Using this sequence fragment, the RdRp gene has been replaced by a sequence encoding the NanoLuc luciferase protein, and the lysis start codon has been removed. Locations of start/stop codons are labelled with green/red bars, and the MinJu long-distance interaction is highlighted in blue.

To demonstrate this, I have constructed a control mRNA sequence containing the MS2 coat and NLuc proteins, CTnano-cnt, by directly replacing the RdRp gene in bacteriophage MS2 with an mRNA sequence encoding the NanoLuc luciferase protein which has been codon optimised for expression in *E. coli* (Genbank ID MN834152). The thermodynamic minimum free energy fold of this mRNA when the MinJu interaction is present (Figure 3a and Supplementary Figure 2) suggests that the TR stem-loop is present, and thus should have similar expression of the nano luciferase protein to that of RdRp in phage MS2. However, when examining the structure of the RBS for the NanoLuc luciferase gene after ribosome disruption of the MinJu sequence, the CTnano-cnt mRNA is predicted by my ribosome/folding kinetics model^17^ to sequester the NLuc start codon into a more stable hairpin during co-translational folding, reducing expression (left hand side of Figure 3b).

**Figure 3.**
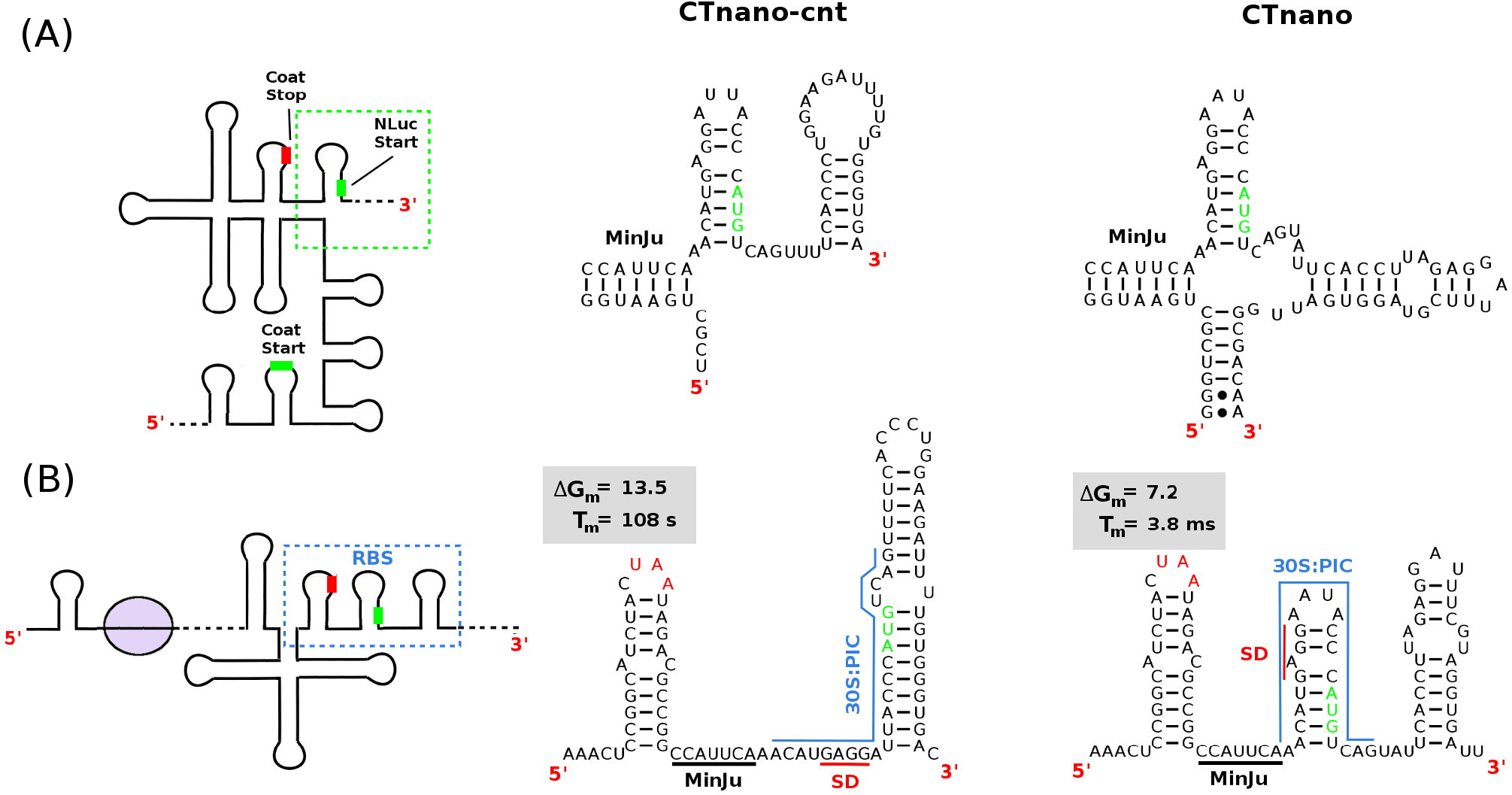
Predicted secondary structures of the NLuc ribosome binding sites in the CTnano-cnt and CTnano mRNAs. (A) Predicted structure of the RBS when the MinJu long distance interaction is present. Ribosome binding to the NLuc RBS should be blocked in both CTnano-cnt and CTnano due to a lack of a ribosome standby site. (B) Predicted structure of the NLuc RBS when the MinJu interaction has been disrupted by ribosomal translation of the coat gene. The estimated energy barrier (Δ*G*_*m*_ - kcal/mol) and the predicted average time (*T*_*m*_) until the binding site for the 30S:PIC (blue line) becomes exposed are shown, along with the Shine-Dalgarno sequence (red line).

In order to prevent formation of this hairpin during co-translational folding, I have synonymous re-coded the NanoLuc luciferase gene in CTnano-cnt mRNA creating the CTnano mRNA (see methods for procedure). The predicted secondary structure of the CTnano mRNA when the MinJu interaction is present/absent (right hand side of Figures 3a and 3b, respectively) shows that the minimum free energy fold now predicts the start codon being sequestered in the TR hairpin, which has a much lower energetic barrier to melting (c.f. Δ*G*_*m*_ in Figure 3b), and thus should result in higher expression levels of the NLuc protein.

### Theoretical prediction of NLuc protein expression

In order to theoretically estimate the levels of NLuc that would be produced from the CTnano-cnt and CTnano mRNAs, one must take into account the competition between ribosomes and coat proteins for the TR stem loop and must also calculate the time dependence on the opening of the ribosome stand-by site for the NLuc gene as a result of the translational coupling between the Coat and NLuc genes. This means that the production rate of NLuc will depend on the production rate of coat protein and the speed of the ribosome movement over the coat gene (the coupling effect), as well as the amount of coat proteins present and their ability to compete with free ribosomes for the TR stem-loop (the repression effect). As discussed in Methods, my computational ribosome/folding kinetics model^17^ is able to account for both the translational coupling as well as the competition between ribosomes and coat proteins for the TR stem-loop. This is since it predicts the co-translational folding of the CTnano mRNA and the rate of movement of the ribosome over the coat gene while also considering the translation of all ribosomes that would be present in the cellular environment, allowing for a calculation of the competition between free ribosomes and coat proteins to be estimated.

Using the secondary structures predicted for the CTnano-cnt and CTnano mRNAs (Supporting Figures 2 and 3, respectively), I simulate the protein expression from multiple copies of either the CTnano-cnt or CTnano mRNA in the presence of the ≈ 1200 background cellular mRNAs and 15000 active ribosomes that would be present in an exponentially growing *E. Coli* cell. The CTnano-cnt and CTnano mRNAs are modelled as being continuously produced with rate *β* from a plasmid vector after induction with IPTG which occurs at time point *t* = 0. The production rate *β* = 0.012 s^−1^ gives the best fit to the experimental data (see below) and results in roughly 20 copies of the mRNA being present in the cell, on average, within 30 min of induction. Figures 4a and 4b show the results of coat protein and NLuc synthesis in both mRNAs over the course of 30 minutes. While the amount of coat protein produced in both cases are roughly identical as expected, the CTnano mRNA is predicted to produce approximately 4.6 times as much nano luciferase as the CTnano-cnt mRNA prior to repression by the coat protein.

**Figure 4.**
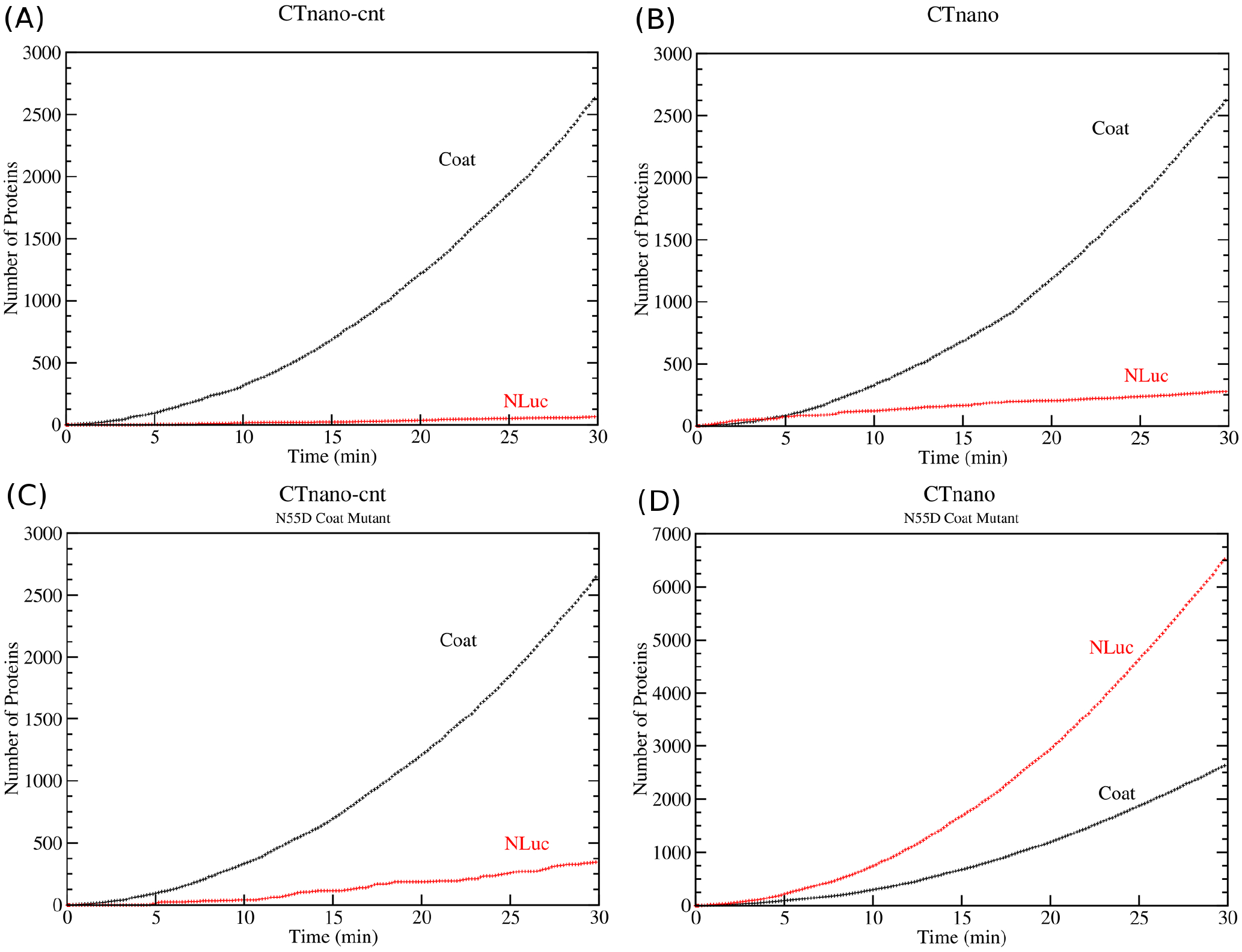
Computational prediction of protein expression in the CTnano-cnt and CTnano mRNAs. Protein expression of the MS2 coat and NanoLuc luciferase genes are simulated using the ribosome/RNA folding kinetic model^17^ assuming multiple copies of the mRNA being produced in the cell at rate *β* = 0.012 s^−1^. Time courses for the amount of MS2 coat protein and NLuc produced for; (A) CTnano-cnt mRNA, (B) CTnano mRNA, (C) CTnano-cnt mRNA with N55D mutation, and (D) CTnano mRNA with N55D mutation.

In addition to the simulation where CTnano and CTnano-cnt mRNAs code for the wild-type MS2 coat protein, I also perform a protein expression simulation of the mRNAs where the MS2 coat protein has the N55D mutation present. This coat protein mutant has been shown to be incapable of binding to the TR stem-loop^27^. Thus, these mutant mRNAs should not be able to repress NLuc synthesis and hence represent the maximum protein production rate that is possible. The computer simulations in Figures 4c and 4d show that the maximum production rate of NLuc in the CTnano mRNA is roughly 20 times that of the CTnano-cnt mRNA, indicating that the re-coded CTnano mRNA has enhanced NLuc protein expression compared with the CTnano-cnt mRNA.

### Experimental measurement of NLuc protein expression

In order to validate the theoretical predictions of NLuc production in the CTnano-cnt and CTnano mRNAs, I have constructed the plasmids pET-CTnano-cnt and pET-CTnano, where expression of the mRNAs are under the control of the T7 promoter (see Methods). Small cultures of *E. coli* cells containing one of the plasmids were grown in LB media and the resulting luminescence from a sample of cell culture was measured with the NanoGlo assay (Premega) as detailed in Methods. The raw data was normalised by the final *A*_600_ of the culture and the data was fitted to a sigmodial function (see supplementary figures 5 and 6). The best fit peak luminescence for both the CTnano and CTnano-cnt mRNAs, along with the peak luminescence from mutant pET-CTnano and pET-CTnano-cnt plasmids containing one of the three coat mutants (N55D, T19*, and S37*), are shown in Figure 5. As can be seen, there is a clear and significant increase in expression of NLuc in the CTnano mRNA when compared with the CTnano-cnt mRNA. Table 1 gives the luminescence / *A*_600_ measurements for each of the mRNAs.

**Figure 5.**
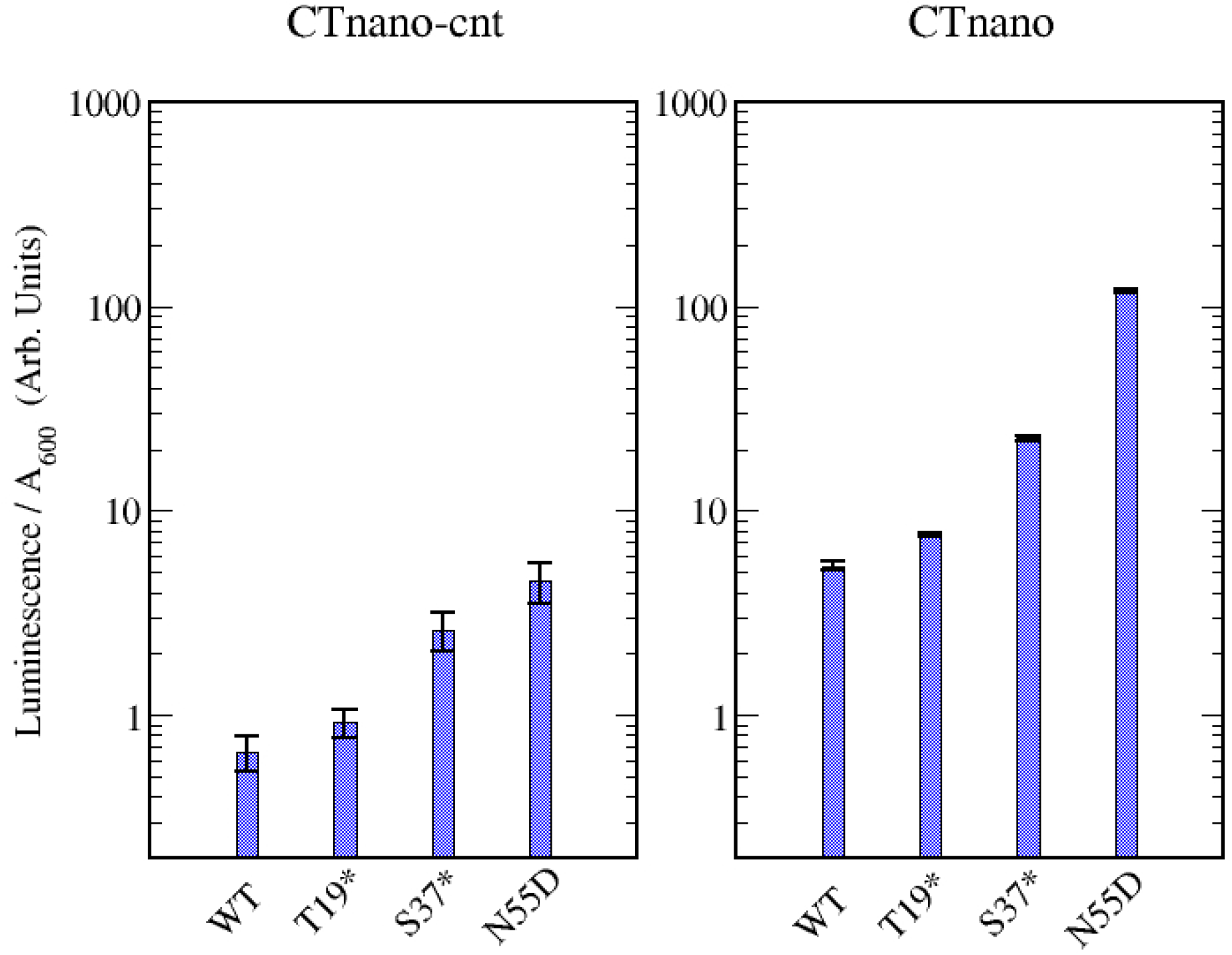
Experimental measurements of peak luminescence produced by the CTnano-cnt and CTnano mRNAs and their mutants. Experimental luminescence values divided by the *A*_600_ of the culture are shown for the CTnano-cnt mRNA (left hand side) and CTnano mRNA (right hand side). Note, experimental data have been divided by 10^4^ for ease of plotting.

**Table 1.** Experimental and theoretical measurements of protein expression in the CTnano-cnt and CTnano mRNAs and their mutants. Experimental measurements have been divided by 10^4^. Theoretical calculations are the average protein numbers over three simulations. Not determined, n.d.

| mRNA | Mutation | Exp.<br>(Arb.U.) | Theory<br>(Num Prot.) |
| --- | --- | --- | --- |
| CTnano-cnt |  | 0.66 | 61 |
| CTnano-cnt | T19* | 0.93 | n.d. |
| CTnano-cnt | S37* | 2.62 | n.d. |
| CTnano-cnt | N55D | 4.57 | 342 |
| CTnano |  | 5.36 | 267 |
| CTnano | T19* | 7.60 | n.d. |
| CTnano | S37* | 22.58 | n.d. |
| CTnano | N55D | 117.67 | 6534 |

#### NLuc is translationally coupled to MS2 coat expression

To demonstrate the theoretically predicted translational coupling in the CTnano and CTnano-cnt mRNAs, nonsense mutations can be introduced to the coat protein before and after the MinJu long distance interaction. It has been previously shown that nonsense mutations introduced prior to the MinJu sequence prevents full expression of the MS2 RdRp protein, while nonsense mutations after increase RdRp expression^26^. Following these experiments that were done to demonstrate translational coupling in phage MS2, I have constructed the plasmids pET-CTnano-M3, which has the nonsense mutation T19* before MinJu and pET-CTnano-M7, which has the alternative nonsense mutation S37* after MinJu. Supporting Figure 4 gives a cartoon diagram of the positions of these stop codons, relative to the MinJu long-distance interaction. Table 1, and Figure 5 show that for both CTnano and CTnano-cnt mRNAs, the T19* mutation results in a similar level of NLuc production compared to wild-type coat protein. Since the 19 amino acid N-terminal fragment of the coat protein that is produced by the T19* mutant is unable to bind to the TR stem-loop, NLuc expression would be expected to be at similar protein expression levels seen in the N55D mutant if the NanoLuc start codon was not coupled to the translation of the coat gene. The translational coupling of NLuc to coat protein is further supported by the S37* mutant, which results in much higher expression of NLuc in both CTnano and CTnano-cnt mRNAs due to disruption of the MinJu sequence. These results are consistent with the NLuc being translationally coupled to the upstream MS2 coat protein.

#### NLuc is translationally repressed by MS2 coat protein

Finally, in order to demonstrate experimentally that NLuc protein production is also repressed by MS2 coat protein in the CTnano and CTnano-cnt mRNAs, I have constructed the plasmid pET-CTnano-M2 which contains the N55D mutation to the coat protein. This coat protein mutant was previously shown to have essentially no affinity for the TR hairpin^27^, and thus will be unable to bind to the TR stem-loop and block initiation of the ribosome on the NLuc start codon. This is clearly demonstrated in the data for both the CTnano-cnt and CTnano mRNAs, which both have a dramatic increase in NLuc production, with a larger increase seen for the CTnano mRNA. Specifically, CTnano-M2 mRNA produces roughly 22 times as much NLuc when compared with CTnano containing the WT coat protein. Likewise, CTnano-cnt-M2 mRNA produces 6.8 times as much NLuc when compared with CTnano-cnt mRNA (c.f. Figure 5 and Table 1). These experimental results are consistent with NLuc production in these mRNAs being suppressed by MS2 coat protein.

### Comparison of theoretical and experimental measurements

The experiment measures the intensity of light produced by the NanoLuc enzyme for a given sample of culture while the theoretical calculations predict number of NLuc proteins produced. To compare with experiment, I assume that NanoLuc enzyme follows Michaelis–Menten kinetics and that light production will be directly proportional to the number of enzymes. Thus, experimental measurements can be compared to theretical ones by examining the ratios. In order to fit the model to the experimental data only two parameters need to be adjusted; the rate of mRNA synthesis from the plasmid (*β*), and the size of the footprint of the 30S pre-initation complex (30S:PIC) on the mRNA during initiation. These are the only parameters which require adjusting in the model as the remaining parameters for ribosome kinetics have been fitted from experimental measurements and for RNA folding from Turner energy parameters^17,20,28^.

It should be noted that it is very difficult to theoretically predict the rate of mRNA production from the plasmid as this will depend on the concentration of T7 polymerase in the cell, the number of copies of the plasmid, and mRNA degradation dynamics, amongst other factors. From a simplified perspective, the value of *β* essentially adjusts the total amount of NLuc produced in a certain time by the N55D mutant, as this mutant will be unable to repress NLuc synthesis. Thus, as *β* increases, there is a corresponding increase in the ratio of NLuc produced by the N55D coat mutant to that of the WT where the amount of NLuc will be “capped” due to repression by the coat protein. I have adjusted *β* such that the ratio of NLuc produced by the N55D mutant to that of the WT roughly matches the experimental observations (c.f. ratios in the bottom row of Figure 6). This results in a value of *β* = 0.012 s^−1^, which corresponds to a theoretical prediction of roughly 20 mRNAs being present in a single cell (on average) 30 min post induction.

**Figure 6.**
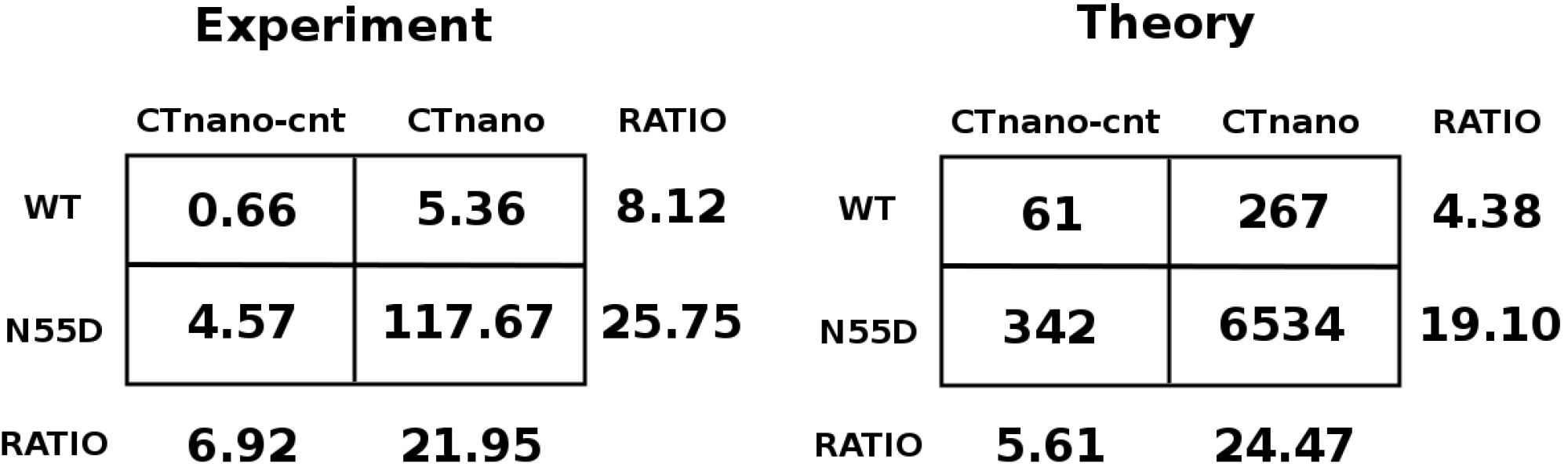
Comparison of experimental and theoretical measurements of NLuc production in CTnano-cnt and CTnano mRNAs. (A) Experimental luminescence / *A*_600_ measurements and (B) theoretical predicted number of NLuc per cell for the CTnano-cnt and CTnano mRNAs containing either the wild-type (WT) MS2 coat protein or N55D coat protein mutant. Ratios between the WT and N55D measurements are shown in the bottom row while ratios between the CTnano-cnt and CTnano mRNAs are shown in the right column. Experimental measurements have been divided by a factor of 10^4^.

The size of the 30S:PIC footprint on the mRNA during initiation determines how much of the RNA needs to become single stranded to expose the 30S:PIC binding site. Thus, if the binding site is sequestered in a more stable RNA secondary structure, the ribosome will take longer to initiate translation resulting in reduced expression of the protein. Again from a simple perspective, altering the footprint will essentially change the amount of the alternative hairpin (present in CTnano-cnt) that requires melting, and will thus impact the ratio of NLuc produced in CTnano-cnt mRNA verses the amount from the CTnano mRNA. I have found that the ideal footprint size corresponds to +6 (−14) nucleotides 3’ (5’) from the A nucleotide in the start codon. This gives the best fit to the experimental NLuc protein ratios in CTnano-cnt verses CTnano mRNAs (c.f. ratios in the right hand column of Figure 6). Interestingly, this footprint corresponds to the amount of mRNA needed to expose an ideally positioned Shine-Dalgarno sequence through to the A site codon. From a comparison to experiment, it is clear that the theoretical melting time for the CTnano-cnt hairpin needs to be slightly longer, or the TR hairpin slightly faster, in order to better match the experimental results. The source of this error is likely due to two possibilities. One possibility is that there could be slight errors in the Turner nearest neighbour model for computing RNA base-pair stacking energies, which would impact on the predicted Δ*G*_*m*_ values and hairpin melting times. Alternatively, there could be inherent errors from using a breadth-first search or greedy algorithm to identify transition paths^17,29^, which are used in the model to compute the estimated energy barriers (i.e. Δ*G*_*m*_) to RNA melting. However, the ability of the model to get the correct qualitative behaviour of the protein expression demonstrates the importance of considering the effects of ribosome movement and competition with other cellular proteins when translational repression is present.

## Discussion

Previous work of David Peabody has demonstrated that MS2 coat protein can be used to repress expression of the RdRp protein on a separate monocystronic mRNA in E coli cells in order to screen for coat protein mutants incapable of binding the TR stem-loop^30^. However, to my knowledge, the possibility of designing a regulatory system on a single polycistronic mRNA in which the MS2 coat protein represses production of a downstream gene other then MS2 RdRp has not been explored. Here I have demonstrated as a proof of concept that such artificial gene regulatory mRNAs can be constructed. Moreover, I have also shown that a stochastic computational model that takes into account detailed steps of ribosome kinetics^17^, can be utilised simulate translational coupling/repression, predict protein synthesis rates, and design mutations which enhance protein expression.

One of the consequences of translational repression mechanisms, such as that observed in bacteriophage MS2 and the synthetic mRNA constructs created here, is that full repression of the gene is delayed due to biding competition between coat dimers and the ribosome for the TR stem-loop which encompasses the start codon. Experimental binding curves from both Peabody and later Lago et al.^27,31^ measure a binding affinity of MS2 coat protein for the TR stem-loop to be around *K*_*d*_ = 1 nM. In the *E. coli* cell, this concentration corresponds to roughly a single coat protein dimer, assuming an average cell volume of roughly 1 fl = 1 *μ*m^3^. Hence one should expect the mRNA to be bound equally as often as it is unbound after synthesis of a single coat protein dimer. However, theoretical calculations reveal that NLuc synthesis continues at the inital synthesis rate until approximately 50 coat protein dimers have been synthesised. This suggests that the TR stem-loop and NLuc RBS is mostly ribosome bound until much higher concentration of coat protein have been achieved to suppress ribosome binding. This highlights the importance of modelling the competition between ribosomes and other cellular proteins for binding to mRNAs when trying to predict protein expression. Translational repression is not unique to phage MS2 and has been observed, for example, in some aminoacyl tRNA synthetases (e.g. metRS thrRS) which have also been postulated to bind to their mRNAs in order to repress protein expression and regulate aminoacyl tRNA synthetase levels in bacteria^32^, and the L11-L1 messenger RNA^16^.

Applying my ribosome/folding kinetic model to CTnano-cnt mRNA translational predicts that co-translational folding of the NLuc RBS after ribosome initiation can result in the formation of two separate hairpin structures, one in which the TR stem-loop is present, and a second more stable hairpin which also sequesters the start codon (c.f. Figure 4 and Supplementary Figure 7). Moreover, the energy barrier to transitioning from the TR stem-loop into the alternative hairpin is predicted to be Δ*G*_*b*_ = 13.8 kcal/mol, while the reverse transition is substantially higher (Δ*G*_*b*_ = 18.1 kcal/mol), suggesting the alternative hairpin is a kinetic trap (see Supplementary Figure 7). Thus, if this alternative hairpin structure is present in the RBS of CTnano-cnt mRNA, coat protein will be unable to bind to this hairpin and the ribosome will continue to express NanoLuc, although at a lower rate due to the higher stability of the hairpin. This also implies that the MS2 coat protein will have lower ability to repress NLuc expression in CTnano-cnt mRNA verses the re-coded CTnano mRNA due to the TR hairpin potentially being re-folded into the alternative hairpin. This is clearly apparent in both the experimental data and theoretical calculations (Figures 5 and 6) which shows that, when wild-type MS2 coat protein is produced by the mRNA, NLuc expression reduces 22 fold in CTnano mRNA compared to the coat mutant N55D. However this reduction is only 7 fold in CTnano-cnt, demonstrating a reduced ability of this mRNA to repress NLuc production. Moreover, the presence of this alternative and more stable hairpin structure reduces the ability of the ribosome to melt the hairpin and access the start codon, resulting in an 8 fold reduction in NLuc expression when wild-type coat protein is produced, or a 25 fold reduction in expression when the N55D coat protein mutant is produced.

Finally, I note that based on the experimental measurements which shows an increasing production of NLuc when nonsense mutations are introduced after the MinJu sequence, it appears that NLuc is translational coupled to the upstream coat gene in both the CTnano-cnt and CTnano mRNAs. As discussed in the Results, if there were no translational coupling, and the ribosome had unrestricted access to the NLuc RBS, then it should be expected that expression of NLuc would be similar to that of the N55D mutant. However, this coupling is slightly leaky as the T19* mutant is still able to express a similar amount of NLuc to that of the wild-type coat protein. A simple explanation for this observation is that the MinJu sequence is able to be disrupted by random thermal fluctuations, given that its melting temperature is on the order of Δ*G*_*m*_ = 8.7 kcal/mol. Currently this can not be simulated in the ribosome folding model as initiation is currently blocked from occurring at internal sites, such as those hidden by the MinJu sequence. Additional work to the model in the future will hopefully enable such features to be simulated. Despite these small needed improvements, my computational ribosome/folding kinetics model^17^ in its current form presents a potentially powerful tool for the recoding of mRNAs to enable both better protein production and the incorporation of advanced features such as built in regulatory mechanisms that control protein synthesis.

## Acknowledgements

I would like to thank Dr. Alex Borodavka, University of Cambridge, and Prof. Alfred Antson, University of York, for graciously providing lab space for me to perform my experiments.

## Author contributions statement

E.C.D. conceived the experiment(s), conducted the experiment(s), performed the computational simulations, analysed the results, and wrote and reviewed the manuscript.

## Data availability statement

Software for computing ribosome translation and mRNA co-translational folding is available on github.com/edykeman/ribofold.

## Additional information

The Author declares no competing interests.

## Notes

### Competing Interest Statement

The authors have declared no competing interest.

## References

1. Rohner, E., Yang, R., Foo, K. S., Goedel, A., and Chien, K. R. (2022) Unlocking the promise of mRNA therapeutics. Nature biotechnology, 40(11), 1586–1600.

2. Qin, S., Tang, X., Chen, Y., Chen, K., Fan, N., Xiao, W., Zheng, Q., Li, G., Teng, Y., Wu, M., et al. (2022) mRNA-based therapeutics: powerful and versatile tools to combat diseases. Signal transduction and targeted therapy, 7(1), 166.

3. Hincer, A., Ahan, R. E., Aras, E., and Seker, U. O. S. (2023) Making the Next Generation of Therapeutics: mRNA Meets Synthetic Biology. ACS Synthetic Biology, 12(9), 2505–2515.

4. Wang, Y.-S., Kumari, M., Chen, G.-H., Hong, M.-H., Yuan, J. P.-Y., Tsai, J.-L., and Wu, H.-C. (2023) mRNA-based vaccines and therapeutics: an in-depth survey of current and upcoming clinical applications. Journal of Biomedical Science, 30(1), 84.

5. Bloom, K., van den Berg, F., and Arbuthnot, P. (2021) Self-amplifying RNA vaccines for infectious diseases. Gene therapy, 28(3-4), 117–129.

6. Papukashvili, D., Rcheulishvili, N., Liu, C., Ji, Y., He, Y., and Wang, P. G. (2022) Self-Amplifying RNA Approach for Protein Replacement Therapy. International Journal of Molecular Sciences, 23(21), 12884.

7. Comes, J. D., Pijlman, G. P., and Hick, T. A. (2023) Rise of the RNA machines–self-amplification in mRNA vaccine design. Trends in Biotechnology,.

8. Green, A. A., Silver, P. A., Collins, J. J., and Yin, P. (2014) Toehold switches: de-novo-designed regulators of gene expression. Cell, 159(4), 925–939.

9. Wang, T. and Simmel, F. C. (2022) Riboswitch-inspired toehold riboregulators for gene regulation in Escherichia coli. Nucleic Acids Research, 50(8), 4784–4798.

10. Karagiannis, P., Fujita, Y., and Saito, H. (2016) RNA-based gene circuits for cell regulation. Proceedings of the Japan Academy, Series B, 92(9), 412–422.

11. Hong, S., Jeong, D., Ryan, J., Foo, M., Tang, X., and Kim, J. (2021) Design and evaluation of synthetic RNA-based incoherent feed-forward loop circuits. Biomolecules, 11(8), 1182.

12. Chappell, J., Westbrook, A., Verosloff, M., and Lucks, J. B. (2017) Computational design of small transcription activating RNAs for versatile and dynamic gene regulation. Nature communications, 8(1), 1051.

13. Chappell, J., Watters, K. E., Takahashi, M. K., and Lucks, J. B. (2015) A renaissance in RNA synthetic biology: new mechanisms, applications and tools for the future. Current opinion in chemical biology, 28, 47–56.

14. Glasscock, C. J., Biggs, B. W., Lazar, J. T., Arnold, J. H., Burdette, L. A., Valdes, A., Kang, M.-K., Tullman-Ercek, D., Tyo, K. E., and Lucks, J. B. (2021) Dynamic control of gene expression with riboregulated switchable feedback promoters. ACS synthetic biology, 10(5), 1199–1213.

15. Peabody, D. (1997) Role of the coat protein-RNA interaction in the life cycle of bacteriophage MS2. Molecular and General Genetics MGG, 254(4), 358–364.

16. Cole, J. R. and Nomura, M. (1986) Translational regulation is responsible for growth-rate-dependent and stringent control of the synthesis of ribosomal proteins L11 and L1 in Escherichia coli.. Proceedings of the National Academy of Sciences, 83(12), 4129–4133.

17. Dykeman, E. C. (2023) Modelling ribosome kinetics and translational control on dynamic mRNA. PLoS Computational Biology, 19(1), e1010870.

18. Groeneveld, H. Secondary structure of bacteriophage MS2 RNA: translational control by kinetics of RNA folding, PhD thesis (1997).

19. Olsthoorn, R. C. L. Structure and evolution of RNA phages, PhD thesis (1996).

20. Mathews, D. H., Sabina, J., Zuker, M., and Turner, D. H. (1999) Expanded sequence dependence of thermodynamic parameters improves prediction of RNA secondary structure. Journal of molecular biology, 288(5), 911–940.

21. Dykeman, E. C. (2015) An implementation of the Gillespie algorithm for RNA kinetics with logarithmic time update. Nucleic acids research, 43(12), 5708–5715.

22. Na, D. and Lee, D. (2010) RBSDesigner: software for designing synthetic ribosome binding sites that yields a desired level of protein expression. Bioinformatics, 26(20), 2633–2634.

23. Salis, H. M., Mirsky, E. A., and Voigt, C. A. (2009) Automated design of synthetic ribosome binding sites to control protein expression. Nature biotechnology, 27(10), 946–950.

24. Salis, H. M. (2011) The ribosome binding site calculator. In Methods in enzymology Vol. 498, pp. 19–42 Elsevier.

25. Seo, S. W., Yang, J.-S., Kim, I., Yang, J., Min, B. E., Kim, S., and Jung, G. Y. (2013) Predictive design of mRNA translation initiation region to control prokaryotic translation efficiency. Metabolic Engineering, 15, 67–74.

26. Berkhout, B. and van Duin, J. (1985) Mechanism of translational coupling between coat protein and replicase genes of RNA bacteriophage MS2. Nucleic acids research, 13(19), 6955–6967.

27. Peabody, D. S. (1993) The RNA binding site of bacteriophage MS2 coat protein.. The EMBO journal, 12(2), 595–600.

28. Dykeman, E. C. (2020) A stochastic model for simulating ribosome kinetics in vivo. PLoS computational biology, 16(2), e1007618.

29. Voss, B., Meyer, C., and Giegerich, R. (2004) Evaluating the predictability of conformational switching in RNA. Bioinformatics, 20(10), 1573–1582.

30. Peabody, D. S. (1990) Translational repression by bacteriophage MS2 coat protein expressed from a plasmid. A system for genetic analysis of a protein-RNA interaction.. Journal of Biological Chemistry, 265(10), 5684–5689.

31. Lago, H., Parrott, A. M., Moss, T., Stonehouse, N. J., and Stockley, P. G. (2001) Probing the kinetics of formation of the bacteriophage MS2 translational operator complex: identification of a protein conformer unable to bind RNA. Journal of molecular biology, 305(5), 1131–1144.

32. Levi, O., Garin, S., and Arava, Y. (2020) RNA mimicry in post-transcriptional regulation by aminoacyl tRNA synthetases. Wiley Interdisciplinary Reviews: RNA, 11(2), e1564.

