## Supplemental Data for "Design of a self-regulating mRNA gene circuit"

### Supplementary Data

**Eric C. Dykeman**

#### Plasmids and primer sequences

DNA sequence constructs containing the CTnano and CTnano-cnt mRNAs used in molecular cloning are given below. The DNA sequences were inserted into the pET-21a(+) vector (Novagen) using restriction sites BgIII and Bpu1102. Supplementary Figure 1 gives a general diagram of the plasmids pET-CTnano and pET-CTnano-cnt.

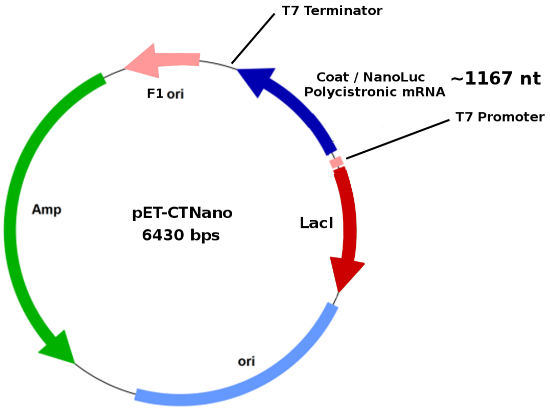


**Sup. Figure 1. Plasmid diagram of pET-CTnano.**

**Primers for Q5 mutagenesis.** Primers and PCR settings used for mutagenesis of the pET-CTnano and pET-CTnano-cnt plasmids are listed in Supplementary Table 1. Amplification of the original plasmid DNA was performed using the Q5 polymerase supplied from the NEB Q5-mutagenesis kit (New England Biolabs) and the primers listed in Sup. Table 1. The PCR reaction was started with denaturation for 1 min at 98° C, followed by twenty-five rounds of PCR amplification. One amplification round consisted of (1) denaturation for 10 sec at 98° C, (2) primer hybridization for 30 sec at $T_a$, (3) polymerase chain extension for 2 min 20 sec at 72° C. The PCR reaction was then finished with a final extension at 72° C for 2 min.

**Sup. Table 1. Primers and PCR settings used in mutagenesis. Primer melting and annealing temperatures are given by the NEB base changer tool (https://nebasechanger.neb.com) in degrees C.**

| Primer Name | Coat Mutation | Primer Sequence | $T_m$ | $T_a$ |
|---|---|---|---|---|
| for-ms2nl-mut2 | N55D | CTCTGCGCAGGATCGCAAATA | 64 | 63 |
| rev-ms2nl-mut2 | N55D | CTCTGACGAACGCTACAG | 62 | 63 |
| for-ms2nl-mut3 | T19* | AACTGGCGACTAGTAAGTCGCCCCAAGCAACTTC | 63 | 64 |
| rev-ms2nl-mut3 | T19* | CCGCCATTGTCGACGAGA | 67 | 64 |
| for-ms2nl-mut7 | S37* | CAGCTCTAACTAGCGTTCACAGG | 65 | 66 |
| rev-ms2nl-mut7 | S37* | ATCCATTCAGCGACCCCG | 68 | 66 |

#### Predicted secondary structures of CTnano-cnt and CTnano mRNAs

Predicted secondary structures of the CTnano-cnt and CTnano mRNAs were computed using standard prediction tools for the thermodynamic minimum free energy. The long-distance MinJu interaction was enforced, assuming that co-transcriptional folding would result in the folding of the coat domain and MinJu interaction by the time the nano luciferase domain was reached by the polymerase. Supplementary Figure 2 shows the structure of CTnano-cnt mRNA, while Sup. Figure 3 shows the structure of the CTnano mRNA along with simplified cartoon diagrams.

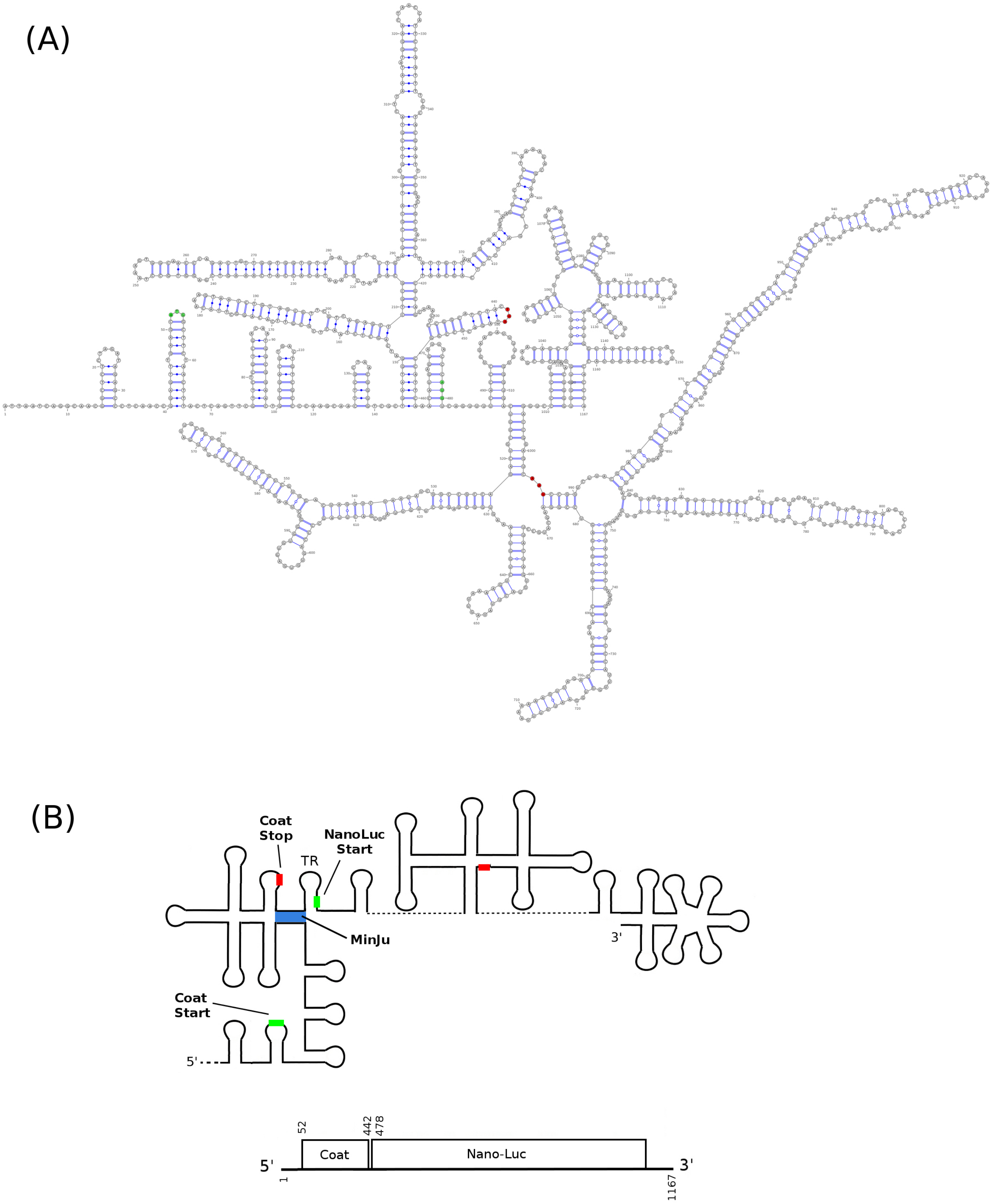


**Sup. Figure 2. Predicted secondary structure of CTnano-cnt mRNA.**

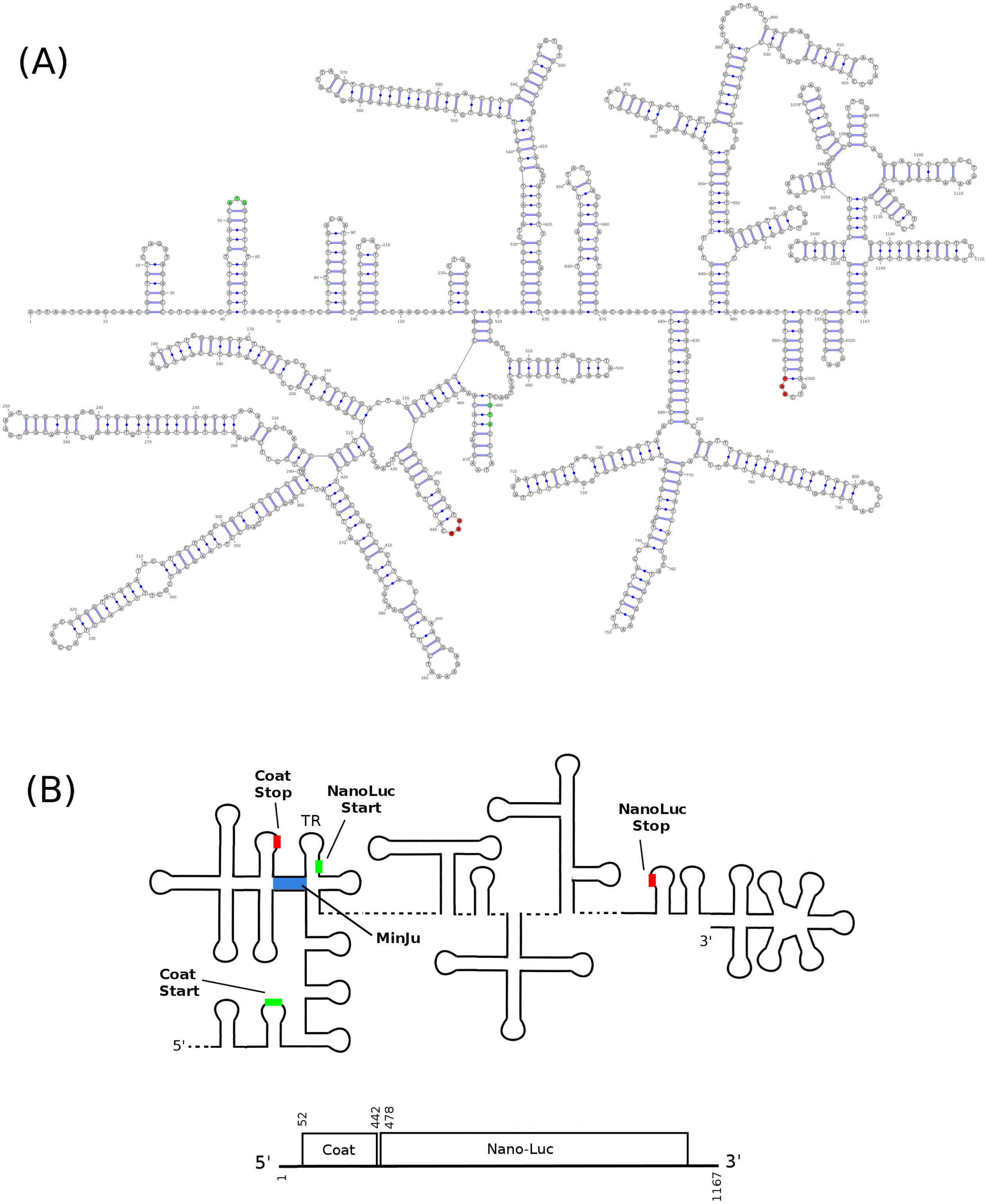


**Sup. Figure 3. Predicted secondary structure of CTnano mRNA.**

#### Stop codon mutations to CTnano-cnt and CTnano mRNAs

Supplementary Figure 4 shows the relative positions of T19* and S37* mutations relative to the position of the MinJu Sequence.

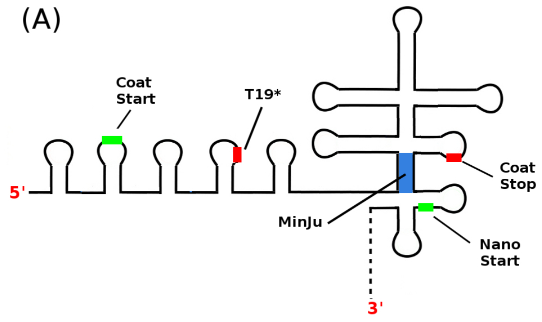


**Sup. Figure 4. The relative positions of T19* and S37* mutations to the position of the MinJu Sequence.** (A) Position of the T19* codon mutation, which is directly upstream of the MinJu long-distance interaction (highligted blue). Ribsomes reaching this stop codon should fail to disrupt the Min-Ju interaction. (B) Position of the S37* codon mutation which is directly downstream of the MinJu long-distance interaction. Ribosomes reaching this stop codon will have fully disrupted the MinJu interaction.

#### Experimental time course of NanoGlo assay

Experimental time courses for the NanoGlo assay are shown for CTnano-cnt mRNA (Sup. Figure 5) and CTnano (Sup. Figure 6) below. Measurements were performed on three culture samples and error bars are the calculated standard deviation of the measurements. Data points were normalised by taking the measured luminescence and dividing by the cultures $A_{600}$ value.

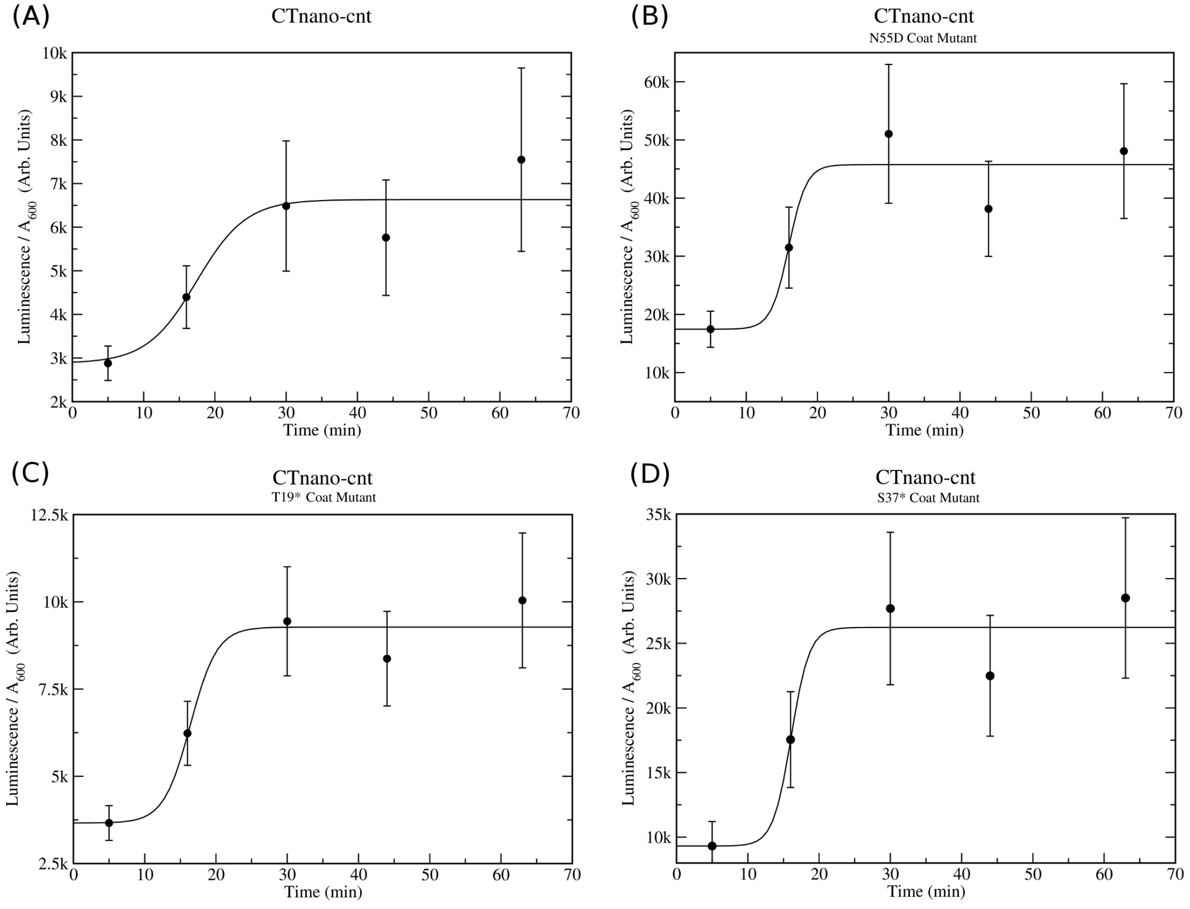


**Sup. Figure 5. Experimental time course of NanoGlo Assay on CTnano-cnt mRNA and its mutants.** Measurements were performed following the procedure in methods and data points are fitted to the equation $L(t) = a + b/(1 + exp(-c(t - d)))$, where $a, b, c$ and $d$ are constants. NanoGlo assays for CTnano-cnt with (A) wild-type MS2 coat protein, (B) N55D coat mutant, (C) T19* coat mutant, and (D) S37* coat mutant.

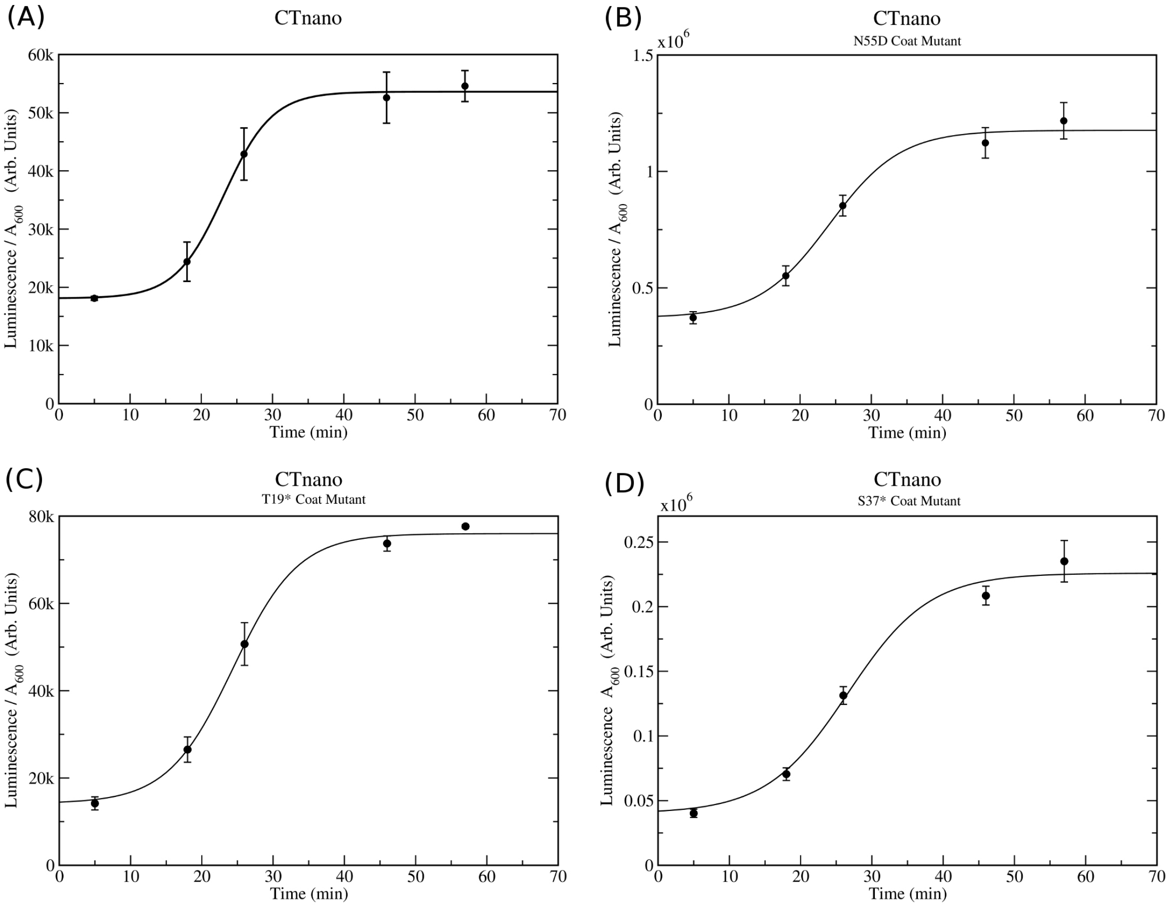


**Sup. Figure 6. Experimental time course of NanoGlo Assay on CTnano mRNA and its mutants.** Measurements were performed following the procedure in methods and data points are fitted to the equation $L(t) = a + b/(1 + exp(-c(t - d)))$, where $a, b, c$ and $d$ are constants. NanoGlo assays for CTnano with (A) wild-type MS2 coat protein, (B) N55D coat mutant, (C) T19* coat mutant, and (D) S37* coat mutant.

#### Co-translational folding kinetics of the CTnano-cnt NLuc ribosome binding site

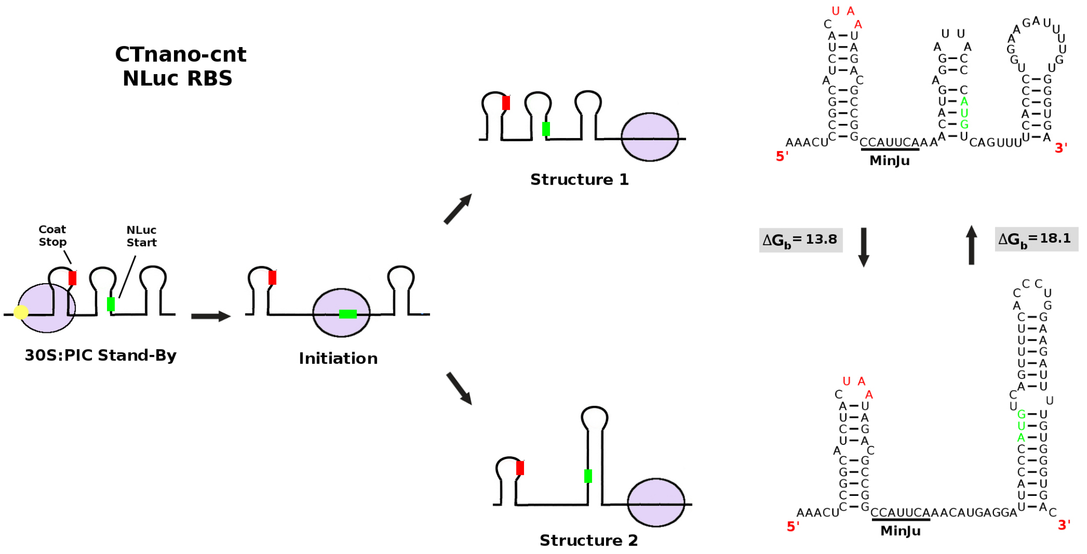


**Sup. Figure 7. Cartoon diagram illustrating the co-translational folding kinetics of the NLuc ribosome binding site (RBS) in the CTnano-cnt mRNA**. Two alternative folding pathways for the RBS can occur as the 5' RNA re-folds during ribosome translation. This results in either the folding of the TR stem-loop (Structure 1), or the folding of an alternative more stable hairpin (Structure 2). The energetic barrier to transition from structure 1 to structure 2 is estimated at $\Delta G_b = 13.8$ kcal/mol. When compared to the barrier between structure 2 and structure 1 ($\Delta G_b = 18.1$ kcal/mol), it is likely structure 2 represents a long lived kinetic trap.

#### Sequences

>CTnano-cnt
AAAGGGAAAGGGAGATCTAATACGACTCACTATAGGGATTAATCAGGCAACGGCTCTCTA
GATAGAGCCCTCAACCGGAGTTTGAAGCATGGCTTCTAACTTTACTCAGTTCGTTCTCGT
CGACAATGGCGGAACTGGCGACGTGACTGTCGCCCCAAGCAACTTCGCTAACGGGGTCGC
TGAATGGATCAGCTCTAACTCGCGTTCACAGGCTTACAAAGTAACCTGTAGCGTTCGTCA
GAGCTCTGCGCAGAATCGCAAATACACCATCAAAGTCGAGGTGCCTAAAGTGGCAACCCA
GACTGTTGGTGGTGTAGAGCTTCCTGTAGCCGCATGGCGTTCGTACTTAAATATGGAACT
AACCATTCCAATTTTCGCTACGAATTCCGACTGCGAGCTTATTGTTAAGGCAATGCAAGG
TCTCCTAAAAGACGGAAACCCGATTCCCTCAGCAATCGCAGCAAACTCCGGCATCTACTA
ATAGACGCCGGCCATTCAAACATGAGGATTACCCATGTCAGTTTTCACCCTGGAAGATTT
TGTGGGTGACTGGCGTCAGACCGCCGGCTATAATCTGGATCAGGTGCTGGAACAGGGTGG
CGTTAGTAGTCTGTTTCAGAATCTGGGTGTGAGTGTGACCCCGATTCAGCGTATTGTTCT
GAGCGGTGAAAATGGTCTGAAAATTGATATTCATGTGATCATTCCGTACGAAGGCCTGAG
TGGTGACCAGATGGGTCAGATTGAAAAAATTTTTAAGGTGGTTTACCCGGTGGATGATCA
TCATTTTAAAGTGATTCTGCATTACGGAACGCTGGTTATTGATGGTGTGACCCCGAATAT
GATTGATTATTTTGGCCGTCCGTATGAAGGTATTGCAGTGTTTGATGGCAAAAAGATTAC
CGTTACCGGAACGCTGTGGAATGGCAATAAGATTATTGATGAACGTCTGATTAACCCGGA
TGGTAGTCTGCTGTTTCGTGTGACCATTAATGGCGTTACCGGTTGGCGTCTGTGCGAACG
CATTCTGGCCTAACTGAGGCCAGGTCCCCCGTAAACGGGGTGGGTGTGCTCGAAAGAGCA
CGGGTCCGCGAAAGCGGTGGCTCCACCGAAAGGTGGGCGGGCTTCGGCCCAGGGACCTCC
CCCTAAAGAGAGGACCCGGGATTCTCCCGATTTGGTAACTAGCTGCTTGGCTAGTTACCA
CCCACTAGCATAACCCCTTGGGGCCTCTAAACGGGTCTTGAGGGGTTTTTTGGTACCAAA
GGCTGAGCAAAGGG
>CTnano
AAAGGGAAAGGGAGATCTAATACGACTCACTATAGGGATTAATCAGGCAACGGCTCTCTA
GATAGAGCCCTCAACCGGAGTTTGAAGCATGGCTTCTAACTTTACTCAGTTCGTTCTCGT
CGACAATGGCGGAACTGGCGACGTGACTGTCGCCCCAAGCAACTTCGCTAACGGGGTCGC
TGAATGGATCAGCTCTAACTCGCGTTCACAGGCTTACAAAGTAACCTGTAGCGTTCGTCA
GAGCTCTGCGCAGAATCGCAAATACACCATCAAAGTCGAGGTGCCTAAAGTGGCAACCCA
GACTGTTGGTGGTGTAGAGCTTCCTGTAGCCGCATGGCGTTCGTACTTAAATATGGAACT
AACCATTCCAATTTTCGCTACGAATTCCGACTGCGAGCTTATTGTTAAGGCAATGCAAGG

TCTCCTAAAAGACGGAAACCCGATTCCCTCAGCAATCGCAGCAAACTCCGGCATCTACTA
ATAGACGCCGGCCATTCAAACATGAGGAATACCCATGTCAGTATTCACCTTAGAGGATTT
CGTAGGTGATTGGCGACAAACCGCCGGCTACAATCTGGATCAGGTGCTGGAACAAGGAGG
CGTTAGCTCCTTGTTCCAGAACCTGGGGGTCAGTGTGACCCCGATCCAGCGTATTGTTCT
GAGCGGTGAAAACGGGTTGAAGATTGATATTCACGTCATCATCCCGTACGAAGGACTCTC
TGGGGACCAAATGGGCCAGATTGAAAAAATTTTCAAGGTGGTCTATCCCGTAGATGACCA
TCACTTTAAAGTGATACTTCACTACGGGACCTTGGTAATCGATGGTGTGACCCCGAACAT
GATCGATTACTTTGGACGGCCCTACGAGGGAATTGCAGTGTTTGATGGCAAAAAGATCAC
CGTTACCGGTACTCTTTGGAACGGCAATAAGATTATTGACGAGCGTCTGATTAACCCAGA
CGGTAGTCTGCTGTTCCGTGTGACCATCAACGGCGTCACCGGTTGGCGCCTCTGCGAACG
AATTCTGGCCTAACTGAGGCCAGGTCCCCCGTAAACGGGGTGGGTGTGCTCGAAAGAGCA
CGGGTCCGCGAAAGCGGTGGCTCCACCGAAAGGTGGGCGGGCTTCGGCCCAGGGACCTCC
CCCTAAAGAGAGGACCCGGGATTCTCCCGATTTGGTAACTAGCTGCTTGGCTAGTTACCA
CCCACTAGCATAACCCCTTGGGGCCTCTAAACGGGTCTTGAGGGGTTTTTTGGTACCAAA
GGCTGAGCAAAGGG
